# Identification of Japanese Encephalitis Virus Genotype V and Other Mosquito-borne Viruses in Camp Humphreys, Republic of Korea, using Metagenomic Analysis

**DOI:** 10.1101/2021.03.15.435489

**Authors:** Mark A. Sanborn, Kathryn McGuckin Wuertz, Heung-Chul Kim, Yu Yang, Tao Li, Simon D. Pollett, Richard G. Jarman, Irina Maljkovic Berry, Terry A. Klein, Jun Hang

**Affiliations:** Viral Diseases Branch, Walter Reed Army Institute for Research, Silver Spring, Maryland, USA; Force Health Protection and Preventive Medicine, MEDDAC-Korea/65th Medical Brigade, Unit 15281, APO AP 96271-5281, USA

## Abstract

Recent outbreaks of emerging and re-emerging viruses such as Zika, West Nile and Japanese encephalitis (JEV) viruses have shown that timely detection of novel arboviruses with epidemic potential is essential to mitigate human health risks. There have been rising concerns that an emergent JEV genotype (genotype V, GV) is circulating in Asia, against which the current US-FDA-approved JEV vaccine may not be efficacious. To ascertain if JEV GV and other arboviruses are circulating in East Asia, we conducted next-generation sequencing on 260 pools of *Culex tritaeniorhynchus* and *Culex bitaeniorhynchus* mosquitoes (6,540 specimens) collected at Camp Humphreys, Republic of Korea (ROK), from mid-May - October 2018. Metagenomic analysis demonstrated a highly abundant and diverse virome with correlates of health and ecological relevance. Additionally, two complete JEV GV genome sequences were obtained from separate mosquito pools, indicating that JEV GV is circulating in the Pyeongtaek area near Seoul, ROK. Retrospective sample and sequence analyses showed that JEV GV was also present in 2016 mosquito pools collected in Seoul, ROK. Sequence-based analysis of JEV GV indicates a divergent genotype that is the most distant from the GIII derived live attenuated SA14-14-2 vaccine strain. A GV E protein investigation and 3D modeling in context to SA14-14-2 indicated likely regions responsible for reduced antibody affinity, including clusters of significant amino acid changes at externally exposed domains. These data highlight the critical need for continued mosquito surveillance as a means of detecting and identifying emerging and re-emerging arboviruses of public health relevance. Importantly, our results emphasize recent concerns that there may be a possible shift in the circulating JEV genotype in East Asia and highlights the critical need for a vaccine proven to be efficacious against this re-emergent virus.

## INTRODUCTION

There has been a dramatic increase in emerging and re-emerging viruses of public health significance, particularly from mosquito-borne arboviruses such as dengue, Zika, West Nile (WNV), chikungunya, and Japanese encephalitis (JEV) viruses (1). Mosquitoes represent a substantial and mostly unexplored virus reservoir of health and ecological relevance. Arboviruses are responsible for roughly 30% of emerging human viruses in recent years (2), and mosquitoes vector plant viruses and are host to insect-specific viruses at extremely high rates (3). The highly pervasive and diverse nature of mosquito viromes makes them hotbeds of virus genetic exchange and evolution and offers observable ecological correlates. Combining routine surveillance with unbiased next-generation sequencing (NGS) of mosquitoes is a highly effective approach for detecting novel viruses and allows for critical insight into the nature of emerging and re-emerging pathogens circulating in vector populations (4–9).

JEV is of great concern due to its severe morbidity and mortality and its continued increase in global distribution. JEV was first isolated in a US service member deployed to Japan in 1935 (10, 11). The geographical distribution now includes 24 countries in Southeast Asia and is the leading cause of tropical viral diseases affecting an estimated 68,000 people per year (12, 13). JEV imposes a high global disease burden (11, 13, 14), with a case fatality of up to 25% of persons demonstrating disease symptoms and with an estimated 50% of persons who survive that exhibit permanent neurological damage, including cognitive dysfunction and neurological deficits (15–19) (13, 20). Although human infections are currently restricted to the eastern hemisphere, JEV genetic material has been identified in mosquitoes and birds in northern Italy, indicating the critical need for surveillance to identify the early emergence of JEV into new geographic locals to prevent the spread of the disease (12).

There are rising concerns regarding a potential genotype shift in the predominant JEV strain circulating in Southeast Asia from genotype I (GI) to genotype V (GV) (5, 6, 21). This is a significant global health concern, as the currently available vaccines have limited reported efficacy against JEV GV (12, 22). JEV GV was first identified in 1952 in Malaysia and was not reported again until it re-emerged 57 years later, where it was detected in *Culex tritaeniorhynchus* mosquitoes from China in 2009 (23). Within the Republic of Korea (ROK), JEV GV was first detected in *Culex bitaeniorhynchus* mosquitoes, collected from Daeseongdong (a village in northern Gyeonggi province located in the demilitarized zone) in 2010, with subsequent detection in *Culex orientalis* and *Culex pipiens* collected in the Gangwon and Gyeonnggi provinces (9, 24). Recently, JEV GV has been reported in clinical cases in the ROK (25).

For this study, we conducted metagenomics-based sequencing of mosquito vectors in the ROK and leveraged our data to make novel virological and ecological insights. By interrogating the data, we uncovered highly abundant and diverse viromes and leveraged statistical analyses to uncover temporal and virus-specific correlations. Additionally, we described our discovery of JEV GV in the context of virome and entomological observations and performed sequence-based analysis that supports the potential for GV vaccine escape.

## MATERIALS AND METHODS

### Sample collection

Mosquitoes were collected using New Jersey light traps or Mosquito Magnets® (Woodstream Corp., Lititz, PA, USA) at Camp Humphreys US Army Garrison. The mosquitoes were identified morphologically using standard keys (26) and pooled (1-39 individuals per pool) by species and collection site and date. The mosquito pools were maintained at -80 °C or on dry ice until processed.

### Viral nucleic acid extraction

Mosquito pools and viral culture media were combined in bead-beating tubes containing glass beads and homogenized using a Mini-BeadBeater 16 (Bio Spec Products Inc., Bartlesville, OK, USA). The homogenates were cleared by centrifugation and the clear supernatants were subjected to nucleic acid digestion with DNase-1, RNase, and Benzonase. The digest was subject to viral nucleic acid extraction using a MagMAX™ Pathogen RNA/DNA Kit on the KingFisher™ Flex Purification System (Thermo Fisher Scientific, Waltham, MA, USA) in a 96 deep-well configuration.

### Random amplification and next-generation sequencing (NGS)

The nucleic acid extracts were treated with DNase-1 prior to a three-step random amplification as described previously (27). Briefly, anchored degenerate octamer primers were annealed with a 65°C to 4°C incubation followed by first-strand synthesis with SuperScript™ III (Thermo Fisher Scientific) and 33 cycles of polymerase chain reaction with Platinum® Taq DNA Polymerase (Thermo Fisher Scientific) and anchor specific primers. Amplicon quality was verified and quantitated prior to library preparation, using the Agilent 4200 TapeStation system and D5000 Screen Tape (Santa Clara, CA, USA). Sequencing libraries were constructed using Nextera® XT DNA Library Preparation Kits and 96 well v2 indexes (Illumina, San Diego, CA, USA). The libraries were quality checked using an Agilent 4200 TapeStation and pooled at equimolar concentrations. Sequencing was performed on an Illumina Miseq system with the 600 cycle v3 Reagent Kit.

### Sequence-based pathogen discovery

The raw paired-end fastq output was processed through an in-house Pathogen Discovery pipeline (28) and the reads were quality evaluated with FastQC (https://www.bioinformatics.babraham.ac.uk/projects/fastqc/) and trimmed using cutadapt v1.16 (29) and prinseq-lite v0.20.3 (30). The quality reads were then assembled into contigs with Ray Meta (31) and extended using Cap3 (32). The contigs then underwent iterative BLAST searching with megablast, discontiguous megablast, and blastx against local NCBI nucleotide (nt) and non-redundant protein databases. The viral sequences assembled and identified by this pipeline were verified using our NGS mapper pipeline (https://github.com/VDBWRAIR/ngs_mapper), which includes data preprocessing followed by reference-based assembly using BWA MEM (https://arxiv.org/abs/1303.3997v2) and outputs assembly statistics and visuals. *De novo* assembled contigs were used as the mapping reference in this pipeline. In addition to screening for assembly errors (e.g., chimeric assembly), sequences published in this study underwent curation involving manual per-base checking of the assemblies for sequencing errors (e.g., incomplete trimming).

### Virome analyses

All viral and unknown contigs assembled by the Pathogen Discovery pipeline were combined with the megablast identified GenBank sequences. These sequences were clustered with CD-HIT-EST (33) at a 90% nt similarity threshold to create a cluster reference sequence set. The cluster reference sequences were then subject to local self-Blastn alignment to identify chimeric sequences and redundancies missed by clustering. For incomplete reference sequences, the total contig file from all samples was mapped iteratively to them to create pseudo-genomes spanning multiple samples (only genomes supported by reads from an individual sample were published in this study). Fragmented or segmented genomes were identified using hierarchal clustering of pool positivity and similarity of Blastx-based identification. BWA MEM was used to map all sample reads to the viral sequence reference set. The resulting alignment files were analyzed with SAMtools (34) and parsed to create a data frame containing sample, sequence, and mapped read data. Correlative hierarchal clustering and Blast identity verification was used to search for redundancies in the data created from segmented viral genomes or incomplete genomes. The resulting master data frame was merged with sample metadata and used for the correlative, prevalence, and other metagenomics analyses. The pooled prevalence rates was calculated with the pooledBin module of the binGroup 2.2-1 R package (35, 36).

### JEV GV confirmation and full genome sequencing

Thirty-five primer pairs were designed for full genome targeted sequencing using the partial JEV GV genome sequence from our unbiased sequencing and the closest GenBank sequence (JF915894) available. The Fluidigm Access Array system (San Francisco, CA, USA) and SuperScript™ III One-Step RT-PCR kit with Platinum™ Taq High Fidelity polymerase (Thermo Fisher Scientific) were used for full genome amplification. Sequencing libraries were prepared with the Nextera® XT kit (Illumina, San Diego, CA, USA) and sequenced on an Illumina Miseq using the v3 600 cycle kit. The genomes were assembled from the read data and curation process as described. The genotypes were assumed by pairwise similarity to other GV published sequences and confirmed by phylogenetic analysis.

### JEV sequence analyses

JEV full genome sequences were obtained from ViPR (https://www.viprbrc.org), screened for obvious errors (e.g., indels), and trimmed to contain only the polyprotein CDS (coding sequence) region. JEV GV E gene sequences were obtained from GenBank or extracted from the full genome sequences. Sequences were aligned by MAFFT (37) and manually checked. The evolutionary models were selected with jModelTest (38) prior to building phylogenies. PhyML (39) with best of NNI or SPR tree space search, estimated base frequencies, and aLRT node support was used to construct full genome phylogenies. FigTree (http://tree.bio.ed.ac.uk/software/figtree/) was used in tree visualization. E gene tree construction, bootstrapping, and visualization were performed using MEGA X (40). Visualizations and genotype comparative analysis were created with custom python coding unless otherwise stated. Pervasive positive selection was tested using HyPhy v.2.0 as described by Pond et al. (41). GV E gene Shannon entropy was calculated using LANL entropy tool (https://www.hiv.lanl.gov/) for the alignment of all the ROK sequences.

## RESULTS

### Unbiased sequencing of mosquitoes captured in ROK reveals diverse virome

The Force Health Protection and Preventive Medicine, MEDDAC-Korea, conducts a nationwide multi-location arthropod-borne disease surveillance program in the ROK (3) (**Figure 1A**). For this study, mosquitoes collected from a single site, Camp Humphreys, Anjeong-ri, Pyeongtaek-si, Gyeonggi province, were analyzed due to its proximity to predominant agricultural areas in Gyeonggi province and the high-density population near Seoul, the capital city (**Figure 1A**). Mosquitoes were collected over 24 weeks from mid-May (05/15/2018) to the end of October (10/31/2018), with up to 40 trap-night collections/week (mean = 31). A total of 78,907 mosquitoes, predominantly comprised of *Culex bitaeniorhynchus* and *Culex tritaeniorhynchus,* both of which are known vectors of JEV. A subset of 6,540 *Cx. bitaeniorhynchus* and *Cx. tritaeniorhynchus* mosquitoes representing collection dates 06/26/2018 to 10/29/2018 were selected and combined into 260 pools by species and date of collection for sequencing. *Culex tritaeniornynchus* mosquitoes made up 163 pools (3,945 specimens) with a median pool size of 30 (range 1-36) and *Cx. bitaeniorhynchus* comprised 97 pools (2,595 specimens) with a median pool size of 30 (range 1-39). cDNA libraries from each pool were sequenced using three MiSeq runs, which produced a total of 1.45×10^8^ paired-end reads for our analysis (**Figure 1B**). *De novo* assembly generated 41,899 contigs with nt lengths ranging from 101 to 15,691 nt, with 1,811 contigs ≥1000 nt (**Figure 1C**). Sequences were aligned to NCBI databases, 1.89×10^7^, 6.40×10^6^, and 1.33×10^5^ reads were classified into super kingdoms *Eukaryota* (non-human), viruses (known and novel), and bacteria, respectively (Figs. 1D-1E). Most viruses were identified as mosquito-specific viruses, plant viruses, unclassified, or uncharacterized viruses with unknown human or animal infectivity. Virus read abundance varied greatly by virus classification and by mosquito species (**Figure 1F**). *Virgaviridae* and *Sobemoviridae*, both known plant pathogens, accounted for the highest number of virus reads found in *Cx. bitaeniorhynchus* and *Cx. tritaeniorhynchus* respectively. Reads classified to known vertebrate pathogen families, e.g., *Orthomyxoviridae*, *Picornaviridae*, *Flaviviridae*, and *Bunyaviridae*, were observed at lesser amounts and in each case were more abundant in *Cx. bitaeniorhynchus*. We also observed reads belonging to mosquito-specific virus families in high amounts, e.g., *Rhabdoviridae* (**Figure 1F**). These data demonstrate that a wide range of potential zoonotic, botanical, and human pathogens can be found in *Cx. bitaeniorhynchus* and *Cx. Tritaeniorhynchus* mosquitoes.

**Fig 1.**
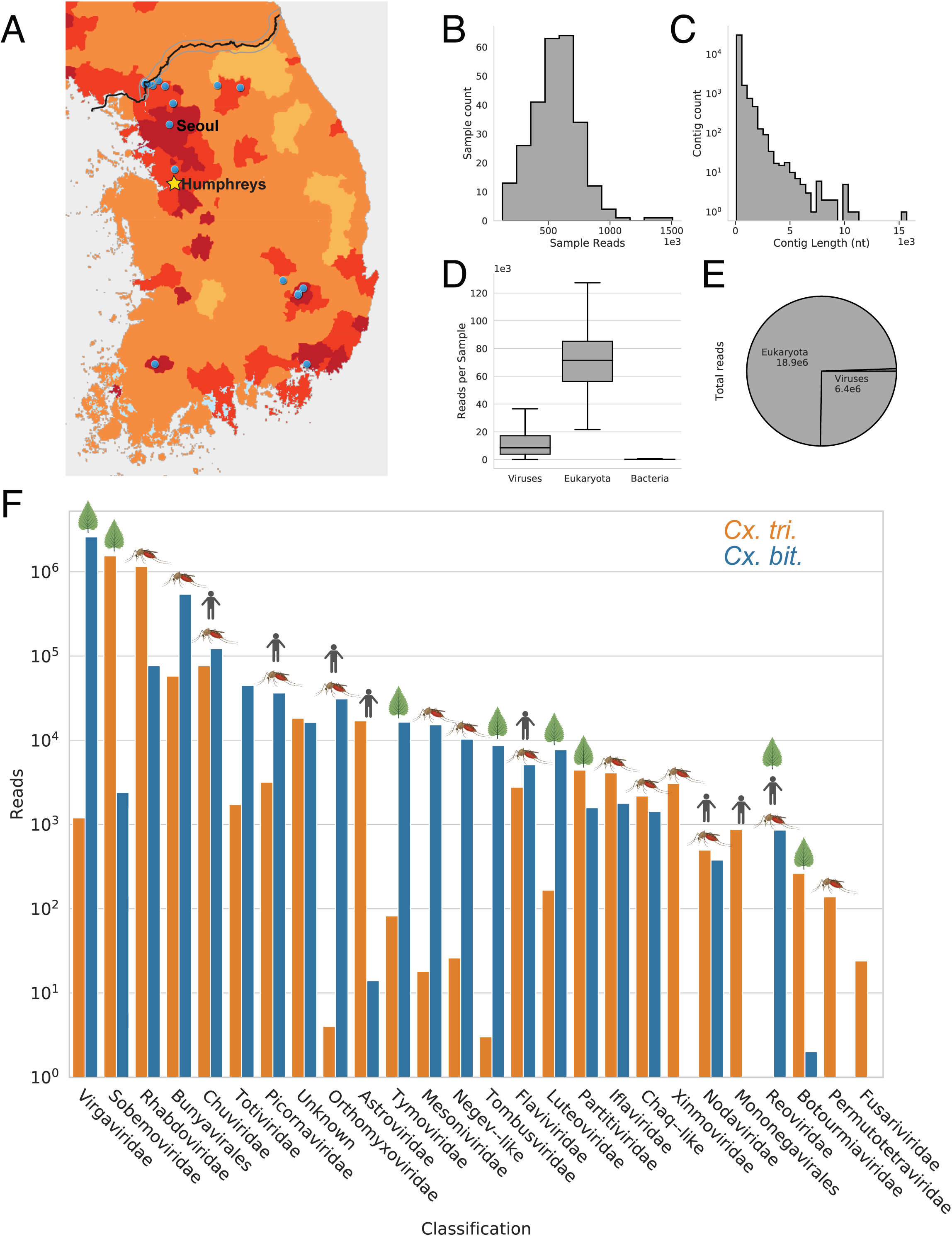
Metagenomics sequence data of the 2018 Camp Humphreys mosquitoes. **(A)** Map of the ROK showing the 2018 Camp Humphreys (star), in addition to other MEDDAC-Korea collection sites (blue dots). Estimated population densities are shown by the shade of orange from dark (high population) to light (low population). (**B)** Distribution of reads per sample. (**C)** *De novo* assembled contig length distribution (**D)** Reads per sample by superkingdom. (**E)** Total reads by superkingdom. (**F)** Plot of total reads by known or putative viral classifications separated by mosquito species. Classifications are ordered from left to right by total number of reads. Known classification tropism (vertebrate, mosquito, plant) is shown by the top cartoons but may not accurately represent many of the highly novel sequences found.

### Viruses are highly abundant in *Culex* populations

We identified 122 unique virus genomes by collapsing all 15,745 virus contigs within 10% nt identity and by identifying segmented and fragmented genomes. The mosquito pools displayed a high rate of virus read positivity with a median rate of 5 genomes/sample (mean = 5.6), while the maximally infected pool contained reads to 17 viruses. A total of 251/260 (96.5%) of the mosquito pools had reads belonging to at least one virus species, and 151 pools contained reads belonging to 5 or more virus species. We then sought to determine the rates at which individual virus species were detected in mosquito populations. Reads of the most prevalent virus, Culex tritaeniorhynchus rhabdovirus, were found in 157 pools. The median number of pools that individual virus genomes were detected in was two (mean = 11.3 samples/genome).

Reads belonging to the 20 most abundant virus genomes were found in 15 or greater mosquito pools (**Table 1**). These virus sequences displayed high individual mosquito infection rates when estimated using binGroup v2.2-1 (35, 36). Sequences belonging to four viruses, Culex tritaeniorhynchus rhabdovirus, Hubei mosquito virus 2, Pyeongtaek Culex Virga-like virus, and Pyeongtaek Culex Bunyavirus, were estimated to have an individual mosquito infection rate of greater than 5%. Culex tritaeniorhynchus rhabdovirus reads, with the highest number of positive pools (n = 157), had an estimated infection of 11.2% in *Cx. tritaeniorhynchus* mosquitoes. Together, these data suggest the population of *Cx. bitaeniorhynchus* and *Cx. tritaeniorhynchus* tested were positive for at least 122 virus species and that these viruses were present at high rates.

**Table 1.** Mosquito infection rates of the 20 most prevalent viruses and JEV GV

| Rank | Novel | Classification | Accession | Name | *Cx. bit.* |  | *Cx. tri.* |  | 3 week maxima |  |
| --- | --- | --- | --- | --- | --- | --- | --- | --- | --- | --- |
|  |  |  |  |  | Positive pools | Infection rate (%) | Positive pools | Infection rate (%) | Infection rate (%) | Month |
| 1 |  | *Rhabdoviridae* | AB604791 | Culex tritaeniorhynchus rhabdovirus | 12 | 0.49 | 145 | 11.18 | > CUL* | NA |
| 2 |  | Sobemo-like | KX882764, KX882765. | Hubei mosquito virus 2 | 7 | 0.28 | 139 | 9.17 | > CUL* | NA |
| 3 | x | Virga-like | MT568541 | Pyeongtaek Culex virga-like virus A18.2454 | 88 | 11.18 | 23 | 0.62 | > CUL* | Aug |
| 4 | x | Bunya-like | MT568528, MT568533, MT568527 | Pyeongtaek Culex bunyavirus A18.3210 | 76 | 6.25 | 17 | 0.46 | 9.4 | July |
| 5 |  | *Bunyavirales* | KM817698, KM817759, KM817727 | Wuhan Mosquito Virus 2 | 0 | 0.00 | 84 | 3.14 | 6.1 | Aug |
| 6 | x | Toti-like |  |  | 61 | 3.78 | 1 | 0.03 | 5.1 | July |
| 7 |  | *Rhabdoviridae* | KU095840 | Tongilchon virus 1 | 60 | 3.69 | 0 | 0.00 | > CUL* | Sep |
| 8 |  | *Flaviviridae* | MG719525 | Quang Binh virus | 1 | 0.04 | 46 | 1.39 | 3.5 | July |
| 9 | x | Sobemo-like |  |  | 0 | 0.00 | 45 | 1.36 | 4.2 | July |
| 10 | x | Orthomyxo-like | MT568529 | Pyeongtaek Culex orthomyxovirus 18-0874 | 34 | 1.59 | 0 | 0.00 | > CUL* | Oct |
| 11 | x | Sobemo-like | MT568530 | Pyeongtaek Culex sobemo-like virus 18-0862 | 0 | 0.00 | 31 | 0.88 | 4.2 | July |
| 12 | x | Unknown |  |  | 27 | 1.22 | 0 | 0.00 | 1.8 | Aug |
| 13 | x | Luteo-like | MT568531 | Pyeongtaek Culex luteo-like virus A18.3206 | 23 | 1.02 | 0 | 0.00 | 3.9 | Sep |
| 14 | x | Sobemo-like |  |  | 0 | 0.00 | 20 | 0.55 | 1.7 | July |
| 15 |  | Sobemo-like | KX882830 | Wenzhou sobemo-like virus 3 | 0 | 0.00 | 19 | 0.51 | 1.7 | July |
| 16 | x | Luteo-like |  |  | 18 | 0.77 | 0 | 0.00 | 3.9 | Sep |
| 17 | x | Sobemo-like |  |  | 0 | 0.00 | 17 | 0.46 | 2.9 | July |
| 18 | x | Sobemo-like |  |  | 0 | 0.00 | 16 | 0.43 | 2.5 | July |
| 19 | x | Sobemo-like |  |  | 0 | 0.00 | 15 | 0.40 | 1.5 | July |
| 20 | x | Sobemo-like | MT568542 | Pyeongtaek Culex sobemo-like virus A18.2268 | 0 | 0.00 | 15 | 0.40 | 3.7 | July |
| NA | x | *Flaviviridae* | MT568538, MT568539, MT568540 | Japanese encephalitis virus | 2 | 0.08 | 0 | 0.00 | 0.5 | NA |

\* All pools were positive in this timeframe and therefore greater than the calculable upper limit (CUL)

### Significant coexistence of viruses

The coexistence of viruses in mosquito populations is an important consideration when predicting possible coinfections or deriving ecological and evolutionary conclusions. We observed significant co-prevalence and correlation of mosquito viruses (Pearson’s r, p ≤ 0.05) in the samples that were visualized using correlative hierarchal clustering and a double-dendrogram cluster map (**Figure 2A**). Because mosquitoes were pooled by collection date, clustering could be used as a proxy for temporal relatedness. For example, several viruses belonging to *Sobemoviridae*, a known plant-infecting family, form a large cluster. Co-prevalence of unrelated virus families was also observed, such as a tight cluster of Chaq-like, Bunyavirales, and Partitiviridae sequences (**Figure 2A**). Significant (p ≤ 0.05) positive partial correlation (i.e., accounting for all variance) was observed for monophyletic virus pairs formed from correlative hierarchal clustering (**Table 2**) and is likely to indicate viruses coexisting in mosquito populations. Moreover, our binary representation of pool positivity allowed us to observe the relative abundance of viruses in the mosquito populations (**Figure 2B**). Relative abundance in combination with distinct hierarchal clustering by species shows that co-prevalence was highly species-dependent. Taken together, these data indicate that the coexistence of viruses is common and even unrelated viruses with differing tropism can be significantly correlated.

**Fig 2.**
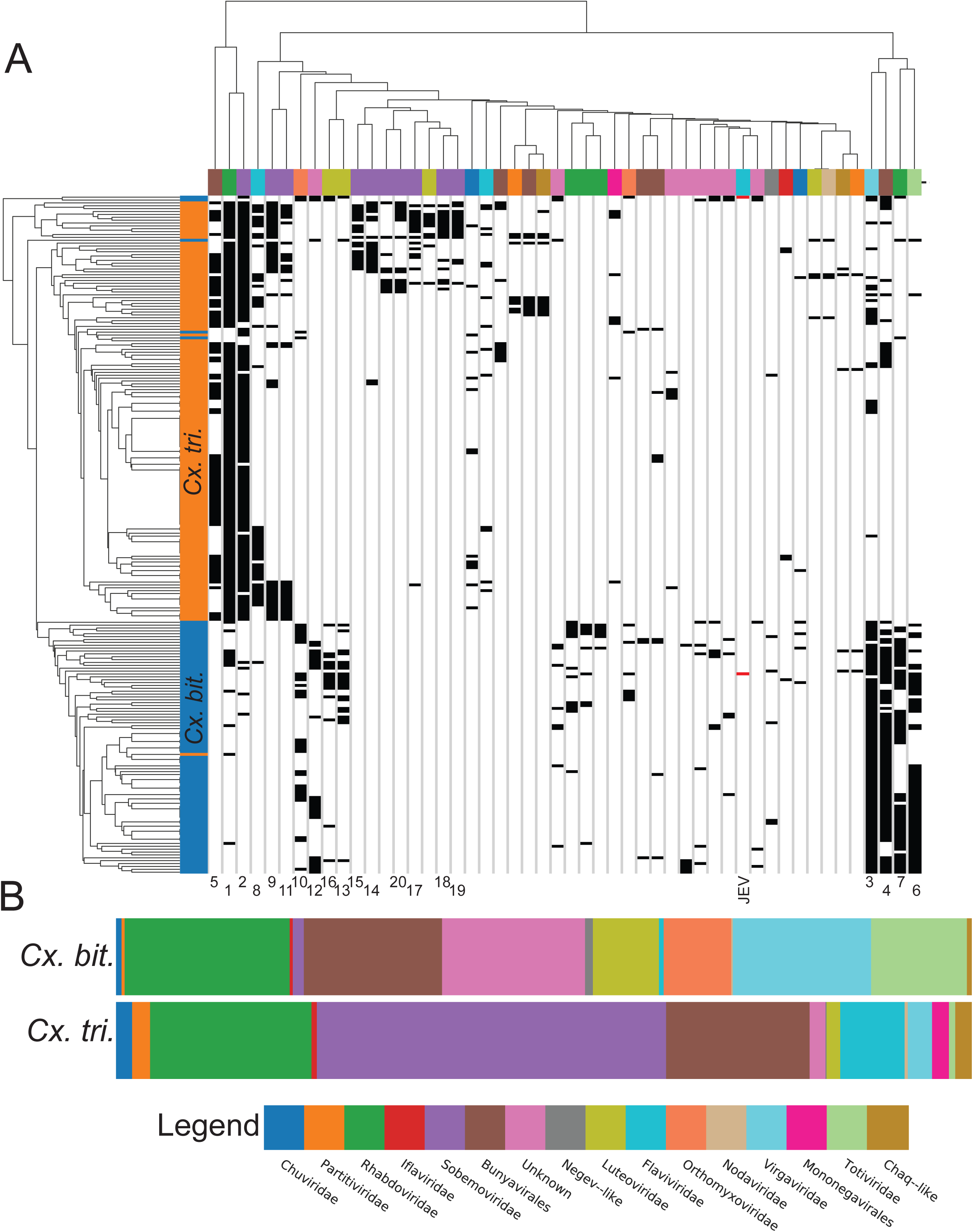
Clustering and relative abundance of viruses found in the mosquito pools. **(A)** Binary clustermap showing the relatedness of unique viruses found in more than 2 pools simultaneously with the relatedness of individual pools with more than 1 identified virus sequence. Virus sequence correlation is depicted by horizontal clustering and the top dendrogram and is color coded (see key) by known or putative classification. Vertical clustering and left dendrogram represents the relatedness of mosquito pools and is color coded by mosquito species. Numbers correspond to rank in Table 1. (**B)** Relative abundance of viruses by mosquito species in the pool as extracted from the clustermap and color-coded according to the key.

**Table 2.** Significantly correlated viruses in mosquito pools

| Virus 1 |  | Virus 2 |  | Partial Correlation |  |
| --- | --- | --- | --- | --- | --- |
| Name | Classification | Name | Classification | r | P value |
| Unnamed | Chaq-like | Unnamed | Partiti-like | 0.85 | 2.20E-35 |
| Unnamed | Sobemo-like | Unnamed | Chaq-like | 0.80 | 3.60E-29 |
| Unnamed | Luteo-like | Pyeongtaek Culex luteo-like virus A18.3206 | Luteo-like | 0.71 | 3.03E-20 |
| Unnamed | Sobemo-like | Pyeongtaek Culex sobemo-like virus A18.2268 | Sobemo-like | 0.70 | 3.02E-19 |
| Wenzhou sobemo-like virus 3 | Sobemo-like | Unnamed | Sobemo-like | 0.63 | 6.27E-15 |
| Unnamed | Sobemo-like | Unnamed | Sobemo-like | 0.56 | 1.91E-11 |
| Pyeongtaek Culex Virga-like virus A18.2454 | Virga-like | Pyeongtaek Culex Bunyavirus A18.3210 | Bunya-like | 0.52 | 4.65E-10 |
| Culex tritaeniorhynchus rhabdovirus | *Rhabdoviridae* | Hubei mosquito virus 2 | Sobemo-like | 0.43 | 4.65E-07 |
| Unnamed | Rhabdo-like | Unnamed | *Rhabdo-like* | 0.43 | 5.98E-07 |
| Tongilchon virus 1 | *Rhabdoviridae* | Unnamed | Toti-like | 0.28 | 1.47E-03 |
| Unnamed | Unknown | Unnamed | Unknown | 0.26 | 3.43E-03 |
| Unnamed | Unknown | Unnamed | Unknown | 0.25 | 4.99E-03 |

Further examination of our data showed that two mosquito pools were positive for JEV, an encephalitic arbovirus of particular significance due to the morbidity and mortality associated with infections. The two individual JEV positive pools were also positive for other viruses. In addition to JEV GV, pool A18.3208 contained sequences belonging to nine additional viruses classified as *Bunyavirales*, *Luteoviridae*, *Orthomyxoviriade*, *Rhavdoviridae*, *Totiviridae*, and *Virgaviridae* (**Sup. Table 1**). Pool A18.3210 had sequences belonging to 14 additional viruses classified as *Bunyavirales*, *Luteoviridae*, *Orthomyxoviridae*, *Rhabdoviridae*, *Virgaviridae*, and unknown classifications (**Sup. Table 1**). These data suggest a wide range of potential co-infecting viruses in JEV positive mosquitoes.

### *Culex* viromes contain temporal and ecological correlates

A complex life cycle exists that supports both transmission and distribution of JEV and other agricultural, zoonotic, and arboviral factors (**Figure 3A**). Our data indicate that viruses have preferential species tropism (**Figure 2A**, **3B**). By projecting the viromes of both *Cx. bitaeniorhynchus* and *Cx. tritaeniorhynchus* pools into Euclidian space (t-SNE), samples distinctively grouped by species (**Figure 3B**). JEV was detected in *Cx. bitaeniorhynchus* pools exclusively, which was surprising as previous studies have reported *Cx. tritaeniorhynchus* as the primary vector of JEV (42). We uncovered seasonal vector-specific prevalence and correlates by binning mosquitoes by collection date and viruses by host tropism. When binned by collection weeks (3-week sliding window), maximum infection rates of the 20 most prevalent viruses were significantly higher and putative plant pathogens peaked temporally in a species-specific manner (**Table 1**). In *Cx. tritaeniorhynchus* pools, eight (8/8) putative plant virus prevalence peaked between late June to mid-July and dropped off by the end of July with small rebounds in September and October (**Sup. Figure 1**). In *Cx. bitaeniorhynchus* pools, three (3/3) putative plant viruses peaked in August, with two (2/3) rebounding again in September corresponding to the detection of JEV GV. In contrast, non-plant viruses showed no temporal trends, suggesting that the prevalence of plant viruses may be more dependent on specific ecological cues, e.g., sap and nectar feeding and seasonal emergence of plant species.

**Fig 3.**
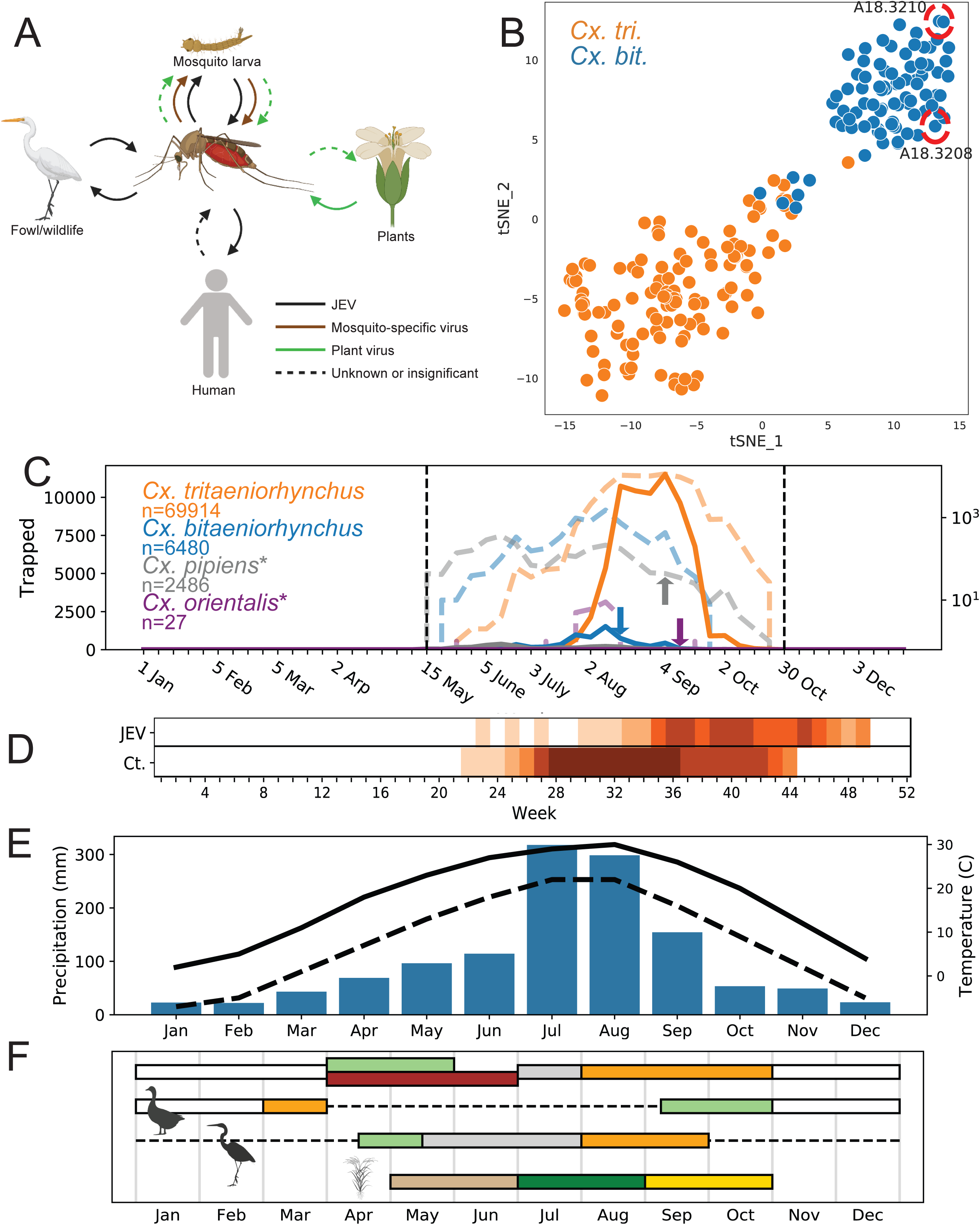
Ecological and temporal factors of JEV in ROK. **(A)** *Culex* mosquitoes vector viruses across kingdoms and phylum and play an important role in the maintenance of JEV in addition to plant, arboviral, and vertebrate viruses. (**B)** tSNE plot of the sequenced mosquito pools as determined by virome composition. The two pools positive for JEV are indicated by the dashed red circles. (**C)** Plot of the total number of individuals from the indicated mosquito species collected at Camp Humphreys in 2018. Vertical dashed lines represent the collection periods and the colored dashed plot lines are the logarithmic representation of the same data. Arrows color coded by species, represent the weeks JEV GV was found. * indicates that JEV was found in that species in 2016 in Seoul. (**D)** Heatmap of weeks with human JEV cases and *Cx. tritaeniorhynchus* observed from 2011-2016 with white being no observations and the darkest shade of orange representing observations all 6 years. (**E)** Average weather in Seoul, ROK as reported by NOAA with the solid line representing temperature highs, dashed line representing temperature lows, and the blue bars representing monthly average precipitation. (**F)** Temporal representation of fowl and the rice growing/harvesting season in the ROK (https://www.birdingkorea.com/, http://www.birdskoreablog.org/?p=18224, http://www.fao.org/giews) (58). From top to bottom the bars represent: typical bird season in ROK, Anas Formosa, Chinese egrets, and the rice growing/harvesting season. Fowl behavior is indicated by white for wintering, light green for arrival of migratory birds, brown for nesting, grey for normal activities, and orange for migratory departure. Dashed lines represent when migratory birds are absent in the ROK. The rice season is coded left to right by planting, growing, and harvesting, respectively.

### Ecological and vector determinants of JEV emergence and transmission

With the implication of ecological and vector impacts to the distribution of viruses in mosquito populations, we sought to determine if JEV detection was observed in a wider range of mosquito species and if that tracked with seasonal correlates. To determine if JEV could be more widely detected in mosquito species across multiple years, we performed a retrospective analysis using NGS data for mosquitoes collected between 2012-2018 to search for additional JEV sequences and determine which species were positive for JEV (**Figure 3C**). Two additional pools were positive for JEV, one each in *Cx. orientalis* and *Cx. pipiens* (**Figure 3C**). No JEV sequences were identified in *Cx. tritaeniorhynchus*. Intriguingly, all four pools (including the two positive pools of *Cx. bitaeniorhynchus*) that were positive for JEV were collected in August and September, regardless of the year that they were collected (**Figure 3C**).

We further explored the relationship between mosquito vectors, weather, agricultural and zoonotic events that may correlate with both JEV incidents in mosquitoes and humans (**Figures 3C**-**3F**). Mosquito collections peaked with a similar timeframe as to when JEV was detected in human populations. *Culex bitaeniorhynchus* mosquito populations peaked at 45 specimens/trap-night during the collection week, beginning the second week of August. *Culex. tritaeniorhynchus* populations peaked at 401 specimens/trap-night two weeks later, starting the fourth week of August (**Figure 3C**). Analysis of data previously reported by Bae et al. (42), who examined the temporal relationship between the peak of *Cx. tritaeniorhynchus* detection in the ROK and reported cases of human JEV, demonstrated a temporal relationship where human JEV infections peak four weeks after peak JEV detection in mosquitoes (**Figure 3D**). The peak collection period of mosquitoes in this study corroborated our own findings. The peak numbers of mosquitoes collected occurred between mid-July and mid-September, with the predominant species captured being *Cx. tritaeniorhynchus* (**Figure 3C**). Since the peak in mosquitoes collected occurring consistently during the same period annually, we examined historical precipitation and temperature data in this region during this same period, as both are key factors in larval habitat and growth and in adult mosquito population abundance. Both precipitation and temperatures reached annual peaks between July and August (**Figure 3E**), just preceding the peak numbers of mosquitoes collected in the region. Moreover, mosquito emergence overlapped with the start of the Fall migration of wading birds, which are known to be common reservoirs of JEV (**Figure 3F**). Human activity also increases the time of year in areas where standing water occurs, e.g., rice paddies and during rice harvesting (http://www.fao.org/giews), which increases the potential for exposure to mosquitoes and subsequent vectored pathogens, e.g., JEV (42). In addition, migratory and domestic waterfowl also have increased exposure to mosquitoes, due to their presence in the rice fields at the same time that JEV mosquito vectors are at their peak (**Figure 3F**). These findings suggest that despite *Cx. tritaeniorhynchus* being reported as the primary vector of JEV in many parts of Southeast Asia, other *Culex* species are likely competent vectors of JEV, adding concern that JEV can be vectored outside of ranges specific to *Cx. tritaeniorhynchus* when reservoir hosts are present. These data demonstrate that a combination of human and ecological factors increases the potential risks of JEV transmission between August and October, increasing the risk of human infections as well as transmission and dispersion of the virus through reservoirs in domestic and migratory wading birds.

### JEV genotype shift in ROK

Early reports have indicated a recent shift in the predominant circulating JEV genotype in the ROK (**Figure 4A**) (5, 6, 21). Initially, the endemic JEV genotype identified in the ROK was Genotype III (GIII) until ∼1990 when it shifted to Genotype I (GI) and then most recently to Genotype V (GV), which was first identified in 2010. Prior to our study, only four full JEV GV genomes were available in GenBank for sequence comparison, collected in 1952 (two nearly identical sequences reported), 2009, and 2015 (**Figure 4B**) (23, 25, 43). We utilized targeted sequencing of our JEV positive pools to assemble three full JEV genomes and we confirmed that they belonged to GV using full CDS phylogenomics (**Figure 4C**). Although all GV sequences fell neatly in the same clade, contemporary sequences have diverged significantly since 1952. The minimum intraclade similarity for all available GV full CDS sequences was 90.3%, with the two most distant sequences being the 2009 Chinese strain and the 1952 Malaysian strain (HM596272). In contrast, the minimum intraclade similarities for GI-GIV full CDS sequences are 94.5%, 95.8%, 94.2%, and 95.3%, respectfully, showing greater distance in the GV clade. Excluding the 1952 GV sequences, the minimum intraclade identity increased to 97.4%. Next, we attempted to leverage the greater availability of GV E gene sequences to determine the drivers of JEV GV emergence in the ROK (**Figure 4D**). Only negative selection was observed (p ≤ 0.05) using both one and two population methods, thus speaking to other drivers of GV emergence beyond E gene selection pressure. We observed diversity among the JEV GV E gene sequences found in the ROK in recent years. For example, two JEV GV strains, which were collected on the same day, in the same vector species, and at the same collection site, were more distantly related than sequences from previous years and sources (**Figure 4D**).

**Fig 4.**
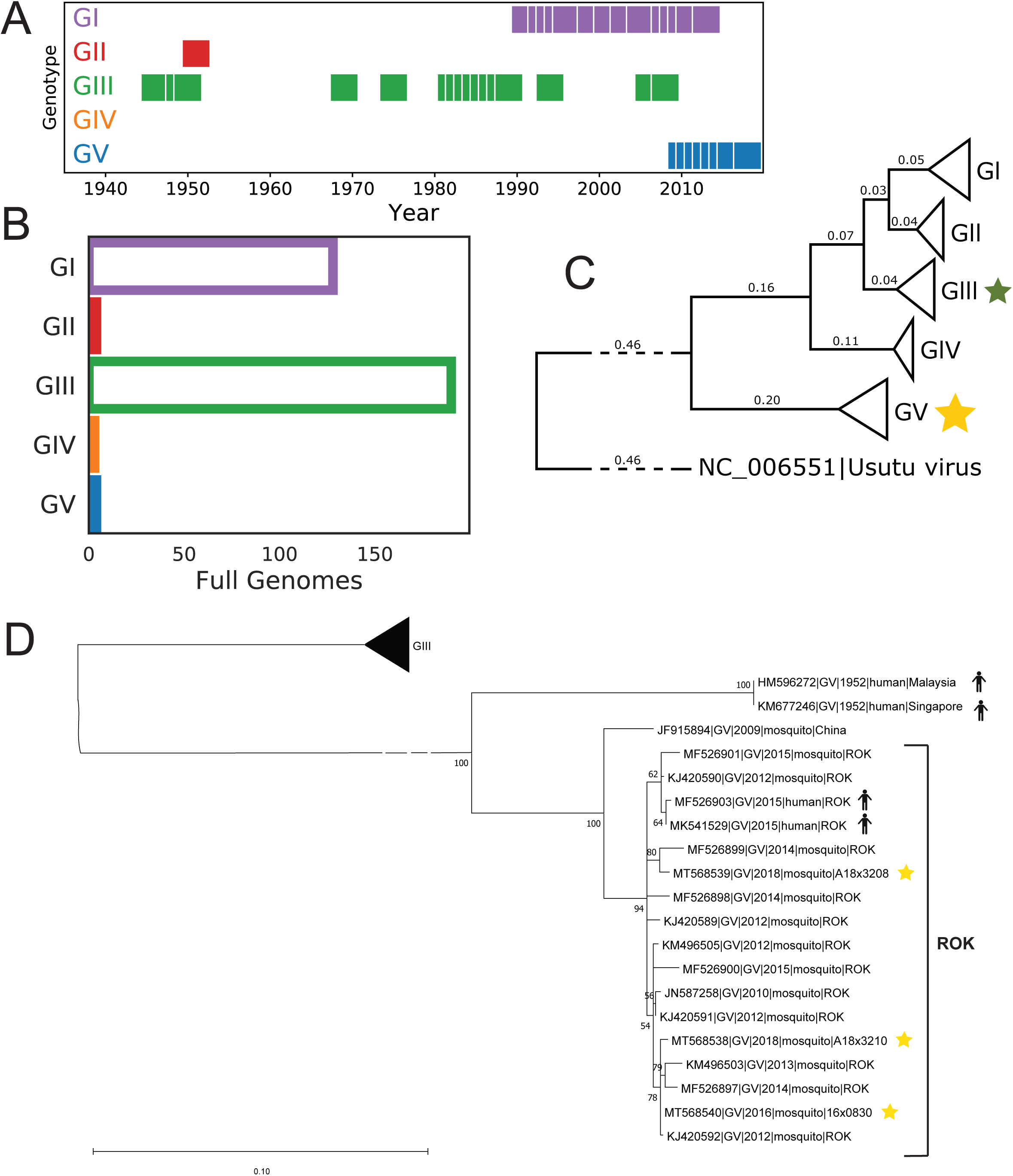
JEV genotype shift in ROK. **(A)** Distribution of JEV genotypes observed in ROK with each square representing a year with an observation. (**B)** Number of full genomes available on genbank by JEV genotype. (**C)** GTR-I-G constructed tree of all available JEV full genomes rooted with an Usutu virus genome. The yellow star represents the clade where the ROK genomes assembled in this study fall and the green star represents the clade containing SA14-14-2 live attenuated JEV vaccine. **(D)** GTR-I-G phylogeny of all available GV E gene nt sequences rooted to a GIII outgroup. Stars denote the sequences found in this study. Bootstrap values are shown.

### Dissimilarity of JEV GV and GIII derived vaccine strain SA14-14-2

The current JEV vaccine strain, SA14-14-2, is based on GIII, raising concerns that the emerging GV may have reduced efficacy due to dissimilarity in key regions of the genome (**Figure 5A**). The average pairwise nt similarities of contemporary circulating GI-GV strains to SA14-14-2 are 88.5, 89.0, 99.2, 84.2, and 78.7 percent, respectively. Sliding window nt similarity analysis (**Figure 5A**) shows a significant drop of similarity between GV and SA14-14-2 in the envelope (E), non-structural (NS) 2a, and NS4b genes. In particular, the E gene that resides on the exposed virion surface is essential for viral entry and contains neutralizing epitopes (44–49). Among JEV GV strains circulating in the ROK, there were observed peaks of increased entropy, suggesting areas of decreased conservation in context to the E gene domains and important immunity motifs (**Figure 5B**).

**Fig 5.**
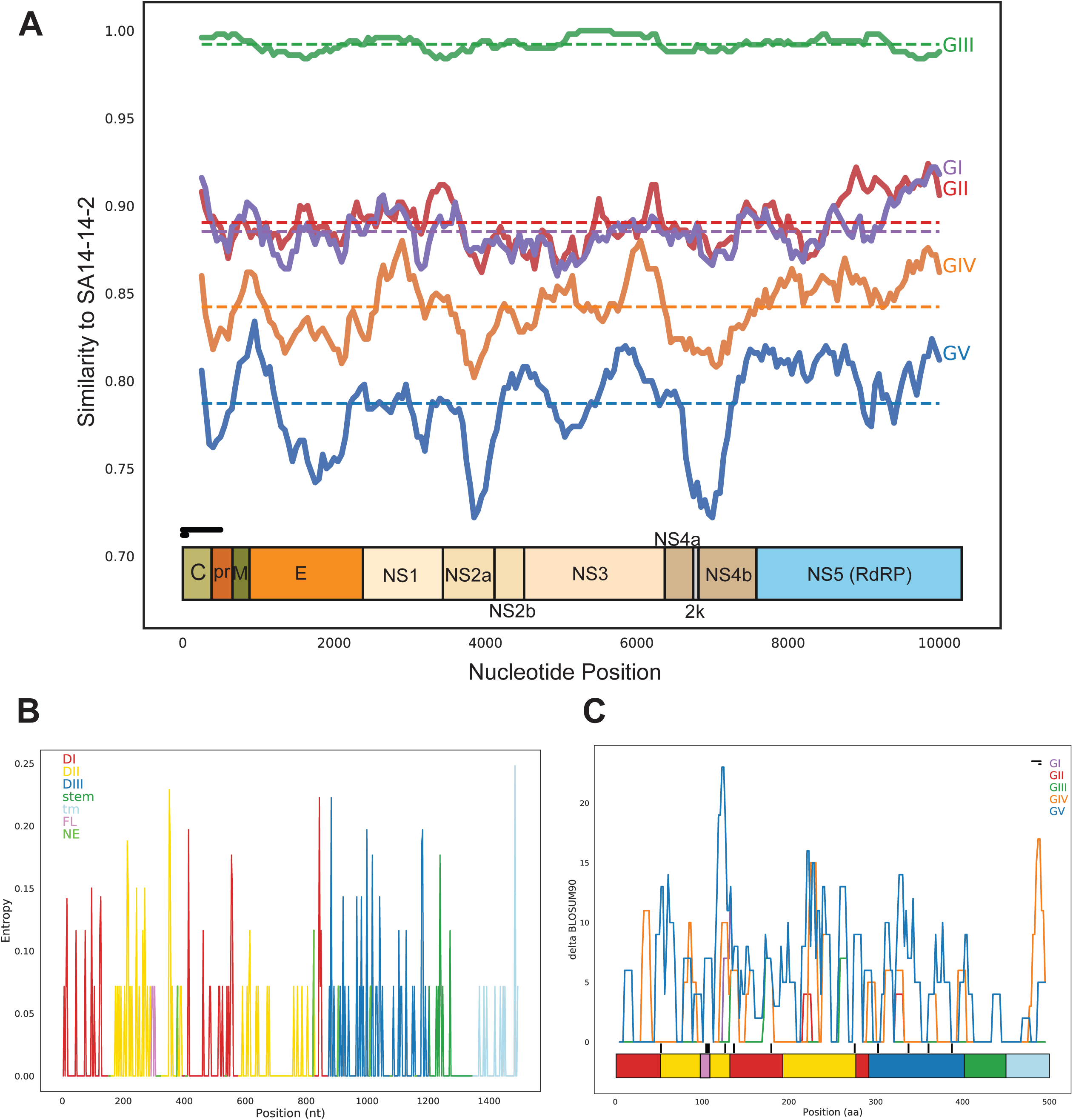
Dissimilarity of JEV GV and GIII derived vaccine strain SA14-14-2. **(A)** Sliding window chart of nucleotide similarity between the natural-occurring JEV genomes collected in or after 2010 (except for GIV which only pre-2010 genomes were available) with SA14-14-2 as baseline. The average identity of each genotype is shown by the dashed lines. The sliding window was 500 nt and the step was 50 nt as shown to scale by the horizontal black bars. The JEV protein layout is shown below. (**B)** Shannon entropy of the 18 contemporary E gene nucleotide sequences. The entropy was averaged by codon. The line chart is color coded by domain. The fusion loop and neutralizing epitopes are shown by the violet and green colors respectively. (**C)** Change in BLOSUM90 aa similarity scores between the E gene consensus aa sequences of each genotype against SA14-14-2 of natural-occurring genomes collected in or after 2010 (except for GIV). The SA14-14-2 E gene tertiary structure domains depicted by the bottom bar and colored by red, DI; yellow, DII; violet, fusion loop; blue, DIII; green, stem; and light-blue, transmembrane region. Vertical black bars represent the position of neutralizing epitopes identified on SA14-14-2. The horizontal black bars represent the window size of 10 and step size of 2 to scale.

We observed areas of increased amino acid (aa) divergence between SA14-14-2 and GV throughout the E protein, most notably a peak directly downstream of the fusion loop, a structure necessary for cell entry (**Figure 5C**). Several other areas of GV and SA14-14-2 aa divergence span known neutralizing, dissimilarity unique to GV. Leveraging the consensus E gene aa sequences of all genotypes, seven GV residues with BLOSUM90 similarity scores less than -1 were identified compared to 2, 2, 2, and 3 for GI through GIV (≤ -1 represents a significant aa substitution). GV and GIV had two aa residue changes (Q52E and I125T) at known neutralizing residues compared to 1 (F107L) present in all genotypes (excluding attenuation sites (50)) (**Figure 6A**). Moreover, by mapping GV and SA-14-2 E protein differences onto the E protein 3D structure,we found a grouping of low similarity aa unique to GV on the exposed D2 domain surface and exposed D3 lateral region (**Figure 6B**). These high-impact residues were in a tight cluster with a handful of lesser-impact differences on the exposed D2 region, and these sites are in close proximity to neutralizing epitopes. Collectively, sequence-based analysis indicates a potential for decreased efficacy of the vaccine strain SA14-14-2 against the emerging JEV GV strain.

**Fig 6.**
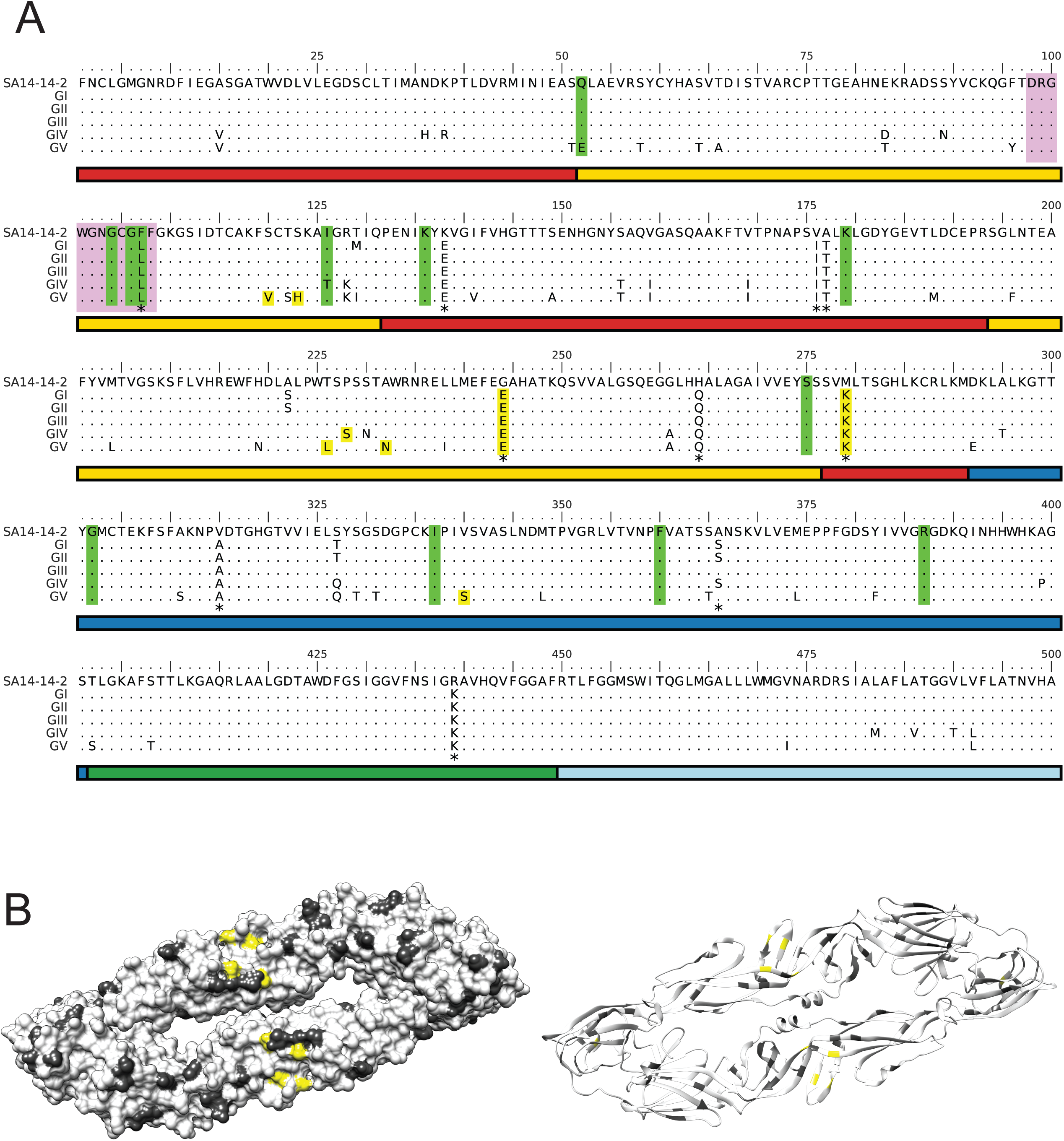
Regions of significant aa divergence in the GV E gene. (**A)** Amino acid alignment of the E gene genotype consensus sequences compared to SA14-14-2. The SA14-14-2 E gene tertiary structure domains depicted by the bottom bar and colored by red, DI; yellow, DII; blue, DIII; green, stem; and light-blue, transmembrane region. The fusion loop is highlighted by the violet box and neutralizing epitopes are highlighted by lime-green. All amino acids with BLOSUM90 scores less than -1 relative to the SA14-14-2 strain are shown highlighted in yellow. * indicates sites of SA14-14-2 attenuation identified by Gromowski et al. (50) (**B)** 3D model of GV changes mapped onto the crystal structure of SA14-14-2 (3p54). All changes are represented by dark grey and unique differences to GV with BLOSUM90 scores of less than -1 are highlighted by yellow. The top exposed surface of the protein is faced towards the reader.

## DISCUSSION

Mosquito surveillance conducted in 2016 and 2018 at Camp Humphreys, near Seoul, ROK, revealed circulation of JEV GV in *Cx. bitaeniorhynchus*, *Cx. orientalis,* and *Cx. pipiens* mosquitoes that are less commonly associated as JEV vectors. The region in and around Seoul has been highlighted in previous studies as a region of key interest in the spread of infectious diseases including JEV, due to the ecological intersect between humans, vectors, reservoirs (e.g., large wading migratory birds) and amplifying hosts (e.g., pigs) (42, 51). Use of metagenomics allowed for an unbiased approach to examine vectored viruses circulating in the ROK that are of human, vector, and agricultural relevance that would have otherwise gone undetected. Metagenomic analysis revealed temporal patterns of JEV emergence in the ROK, corresponding to agricultural and ecological shifts in vector and reservoir habitats. Further characterization of the JEV strains isolated revealed that the genotype detected was GV, both in 2016 and 2018. This is of critical importance, as previous reports have demonstrated that the currently approved vaccine may have limited efficacy against JEV GV (22). This study provides support for concerns regarding a probable shift in the predominant JEV genotype circulating in Southeast Asia toward a genotype for which the vaccine may provide less protection due to inherent sequence differences. Moreover, it highlights the significance of performing routine metagenomics-based analysis based on vector surveillance to detect circulating viruses of global health importance.

Analysis of the vector-specific viromes revealed temporal and seasonal associations of interest. Distinct temporal prevalence was observed in plant pathogens that were not found in other non-plant-associated viruses. Plant feeding patterns and the behavior of mosquitoes, in combination with flora seasonality, may bring them into proximity with competent migratory bird reservoir hosts or other vertebrate hosts, thus increasing the potential transmission of viruses from vector to reservoir as well as typically dead-end hosts (e.g., humans). Moreover, while there is a well-defined role of arthropods in transmitting pathogens of global health relevance, the burden of mosquitoes on agriculture is not a topic well explored despite the abundance of unclassified sequences related to plant viruses identified in mosquitoes in recent years. These plant virus observations in mosquito endemic areas provide data to fill knowledge gaps in virus identification, function, and evolution, and expand our understanding of these health-relevant viromes.

Metagenomic analysis revealed distinct differences in the viromes represented by *Cx. tritaeniorhynchus* and *Cx. bitaeniorhynchus* mosquito pools. Significant co-prevalence of viral sequences was observed in mosquito pools, suggesting that virome composition may contain predictive correlates for health-relevant viruses such as JEV. The distinct temporal and ecological patterns observed in the viruses identified may allow for developing a computational machine learning model to facilitate transmission and disease risk prediction. Models can be refined from continual data input from routine surveillance. This would be of critical epidemiological relevance and provide insights on how to predict and combat future outbreaks of arthropod-vectored viruses.

The high infection rate of the viruses reported here and elsewhere (3, 52, 53) lends importance to their role as models for virus ecology, virus-virus interactions in co-infected hosts/vectors, and virus evolution. In addition to predictive correlates, the highly abundant nature of viral genetic material found in the mosquito pools has implications on virus evolution due to the increased propensity for genetic exchange, especially among like species. Recombination requires coinfection and is common between invertebrate viruses (54). This creates the potential for recombination events within vectors and may contribute increased emerging pathogens worldwide. The correlations between observed sequences hint at viruses and virus families more likely to cause coinfection and possible recombination. From a public health viewpoint, vector-mediated recombination of viruses is of great concern (55). The wide diversity and novelty of viruses discovered by these studies help to fill knowledge gaps in virus evolution and spread. Our observations suggest that important insights would otherwise be missed with more targeted sequencing methodologies. Moreover, these studies are more in line with the One Health approach, emphasizing the importance of multifaceted ecological consideration for the control of zoonotic disease (https://www.who.int/news-room/q-a-detail/one-health).

Previous studies have implicated the co-location of reservoir species including migratory birds with the emergence of JEV (8, 56). Notably, the autumn migratory season occurs during the same period when there are peak mosquito populations and is followed by human JEV cases (51, 57). The west coast of ROK near Seoul has long been identified as a key byway and nesting area for spring and fall migratory routes along the East Asian–Australasian Flyway (EAAF) (https://www.eaaflyway.net/) (58). It is estimated that only ∼10% of birds are native to the ROK, while the remaining species are migratory birds, either spending summers (spring migrants such as Chinese egrets) or winters (such as the Baikal Teal) in the ROK (https://www.birdingkorea.com/, http://www.birdskoreablog.org/?p=18224) (51). These temporal relations reveal a critical time to survey mosquito populations for circulating viruses of human relevance and highlight the potential risk for the spread of JEV from the ROK along migratory routes in the EAAF (51, 59). Our data suggest that JEV and vector emergence overlap with bird migration, which may increase the risk of JEV spread through migratory routes. Our data, along with others (24), also suggest that other *Culex* species are competent JEV GV vectors and add concerns for the transmission of JEV GV outside of ranges specific to *Cx. tritaeniorhynchus*. With the recent shift to GV (42), this poses a significant public health concern due to possible reduced efficacy of current JEV vaccines (22).

Examination of JEV publications demonstrated a potential shift in the dominant genotype in ROK to GV (22, 42). Existing JEV vaccines have an unverified level of protection to GV since published reports raise serious concerns of reduced vaccine efficacy. However, this concern is speculative due to a paucity of contemporary GV isolates and the lack of extensive in-vivo data (22). GV is the most distantly related genotype to the GIII derived vaccine strain and is shown to be more divergent than GI, GII, and GIV. Our analysis showed significant differences in the E, NS2a, and NS4b regions of GV compared to the SA14-14-2 vaccine. The E gene, in particular, is important for cell entry and antibody neutralization. Our data showed that GV has significant aa residue differences from SA14-14-2 at important exposed regions of the E gene. These data hint at possible mechanisms of reduced efficacy, but further in vitro study is required to confirm these observations.

This study revealed the diverse virome associated with *Culex* mosquitoes and identified ecological correlates to JEV. The JEV GV genomes we have identified and sequenced in this study and those in previous studies show significant sequence divergence from that of the current vaccine, reiterating concerns of vaccine efficacy and a shift of the predominant JEV genotype in the ROK. These results demonstrate the essential role of unbiased sequence-based analysis of arboviral vectors in global surveillance programs to characterize and detect emerging and re-emerging pathogens, as well as those of ecological and environmental significance.

## Data availability

Next-generation sequencing raw read data has been deposited under NCBI BioProject ID PRJNA688920. Assembled viral genome sequences were deposited under NCBI GenBank accession numbers MT568527 - MT568542.

## Funding

Funding was provided by the Armed Forces Health Surveillance Division, Global Emerging Infections Surveillance (GEIS) Section, ProMIS ID P0039_18_ME.

## Acknowledgments

We thank Mr. James S. Hilaire, Ms. Nicole R. Nicholas, Mr. Tuan K. Nguyen and Ms. April N. Griggs for their assistance in project management, sample tracking, storage and retrieval. We thank Ms. Tessa Nixon and Ms. Elaina Justiniano for their assistance in data quality control.

## Conflicts of Interest

The authors declare no conflict of interest.

## Disclaimer

Material has been reviewed by the authors’ respective institutions. There is no objection to its presentation and/or publication.

The views expressed in this article are those of the authors and do not necessarily reflect the official policy or position of the Department of the Army, Department of Defense, or the U.S. Government. Authors, as employees of the U.S. Government (KMW, HCK, RGJ, TAK, JH), conducted the work as part of their official duties. Title 17 U.S.C. §105 provides that ‘Copyright protection under this title is not available for any work of the United States Government.’ Title 17 U.S.C. §101 defines a U.S. Government work is a work prepared by an employee of the U.S. Government as part of the person’s official duties.

---

**Supplemental Fig 1.**
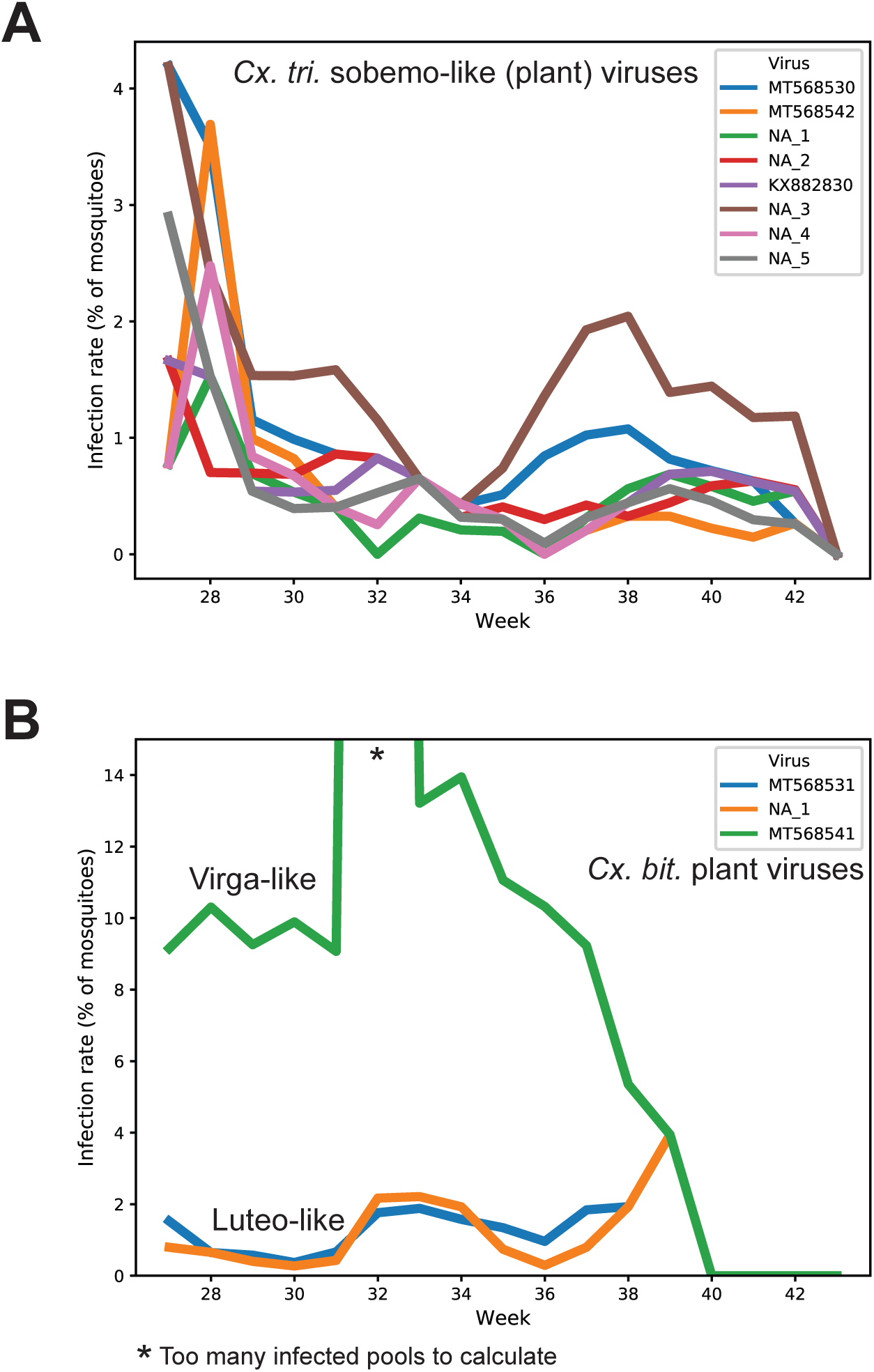
Temporal dynamics of plant viruses in mosquitoes. (**A)** Timeseries of the most highly prevalent *Cx. tritaeniorhynchus* plant viruses (all sobemo-like). Weeks correspond to ISO week dates (27 = late June/early July). (**B)** Timeseries of the most highly prevalent *Cx. bitaeniorhynchus* plant viruses

**Supplemental Table 1.**
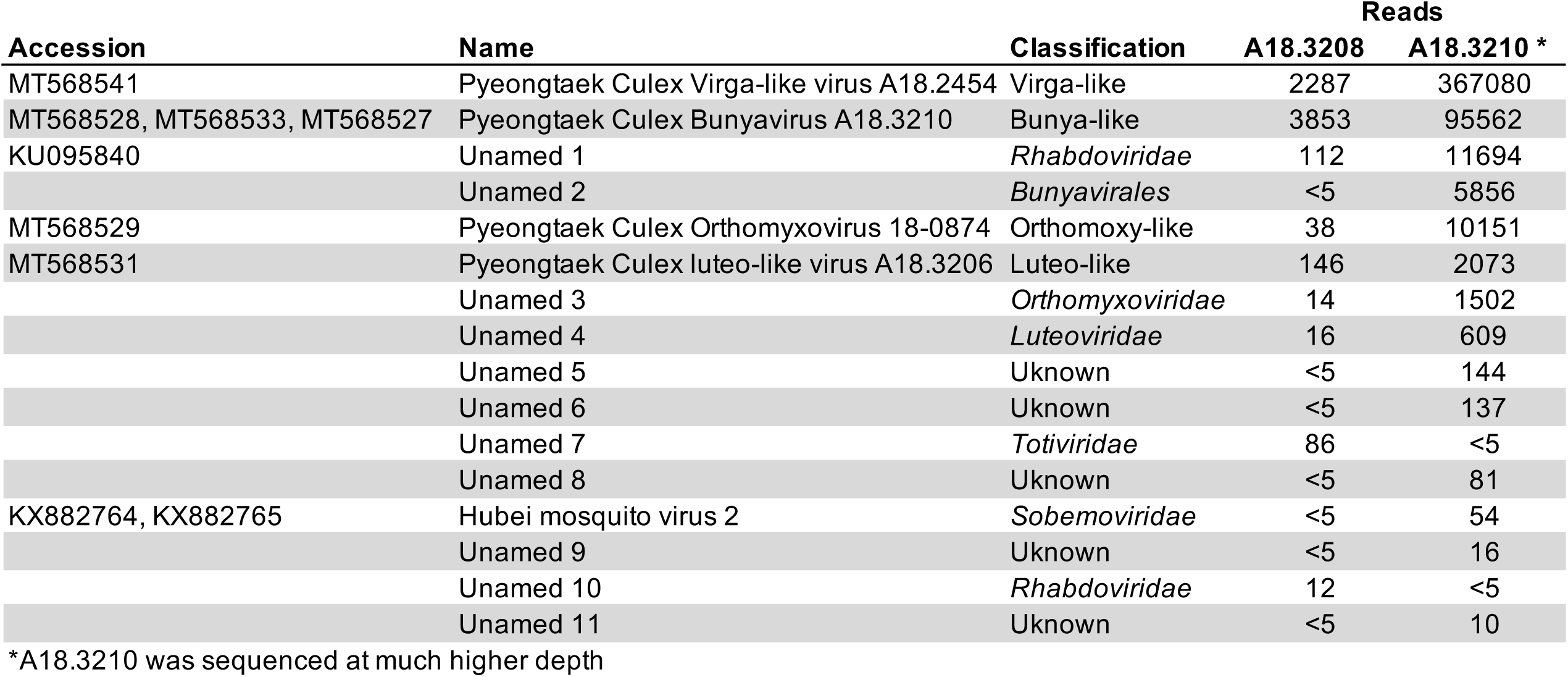
Additional virus reads in JEV GV positive pools

## Notes

### Competing Interest Statement

The authors have declared no competing interest.

